# Microstructural organization of Superior Longitudinal Fasciculus and Cingulum Bundle support metacognition driven cognitive offloading

**DOI:** 10.1101/2024.08.24.609527

**Authors:** Yunxuan Zheng, Bin Bo, Danni Wang, Yiyang Liu, Sam J. Gilbert, Yao Li, Sze Chai Kwok

## Abstract

People often use external tools to offload cognitive demands associated with remembering future intentions. While previous research has established a causal role of metacognition in cognitive offloading, the neural mechanisms supporting this metacognitive control process remain unclear. To address this, we conducted a study with 34 participants using diffusion tensor imaging (DTI) to examine how white matter connectivity supports metacognition-driven cognitive offloading. Behaviorally, we replicated prior findings showing that under-confidence in internal memory predicts a bias toward using external reminders. At the neural level, we used diffusion tensor imaging to quantify fractional anisotropy (FA), a measure of microstructural integrity in white matter. We found the microstructural integrity of the superior longitudinal fasciculus (SLF) and cingulum bundle (CB) negatively predicted deviations from the optimal use of reminders. The microstructural integrity of the fornix negatively predicted participants’ confidence in performing the task when restricted to internal memory. Our findings reveal the microstructural organization of these fronto-temporal-parietal white-matter tracts are related to metacognition driven cognitive offloading. We discuss several aspects of metacognition driven cognitive offloading from a white matter microstructural perspective.

## Introduction

Prospective memory refers to the ability to remember an intention to be fulfilled under an appropriate context in the future (Burgess et al., 2001; Rummel & Kvavilashvili, 2023; Simons et al., 2006). Although successful fulfillment of delayed intentions constitute a substantial part of meaningful life, 50%-70% of memory failures stem from failures in prospective memory (Crovitz & Daniel, 1984). Therefore, many people choose to use external aids to support internal memorization of delayed intentions. For example, a person might write a shopping list before going to the grocery store. In cognitive science, such a strategy of using physical actions to reduce information processing requirements and cognitive demands is known as cognitive offloading (Gilbert et al., 2023; Risko & Gilbert, 2016).

Given the ubiquity of cognitive offloading in everyday life, recent studies have tried to understand when and why people adapt the cognitive offloading strategy (Gilbert, 2024; Gilbert et al., 2023). For instance, cognitive offloading can be considered as a preference to avoid the effort required to maintain information in prospective memory (Chiu & Gilbert, 2024; Dupont et al., 2023; Kelly & Risko, 2022; Sachdeva & Gilbert, 2020). Consequently, memory for offloaded items tends to be relatively poor, if reminders are unexpectedly removed (Eskritt & Ma, 2014; Kelly & Risko, 2019; Sparrow et al., 2011). Recent evidence has also demonstrated that cognitive offloading is causally driven by metacognition, or rather specifically metamemory, the ability to monitor and control one’s own memory process (Dunn & Risko, 2016; Gilbert, 2015; Gilbert et al., 2020; Hu et al., 2019; Risko & Dunn, 2015). That is, people are more likely to commit to cognitive offloading behaviors when they are under-confident about their prospective memory. In contrast, when people feel more confident in their task performance, they would be willing to exert more effort and invest more in their cognitive control (Boldt et al., 2019; Frömer et al., 2021).

However, little is known about how the decision to use external aids and offload memory demand is supported in the brain at the white matter microstructural level. Previous functional neuroimaging research has highlighted the role of the rostral prefrontal cortex (rPFC) in maintaining delayed intentions, with the medial and lateral rPFC serving dissociable functions in this process (Burgess et al., 2001, 2003; Gilbert, 2011; Momennejad & Haynes, 2013; Simons et al., 2006). Specifically, the medial rPFC has been hypothesized to maintain the specific details of delayed intentions, while the lateral rPFC plays a more content-free role (Landsiedel & Gilbert, 2015). A recent study by Boldt and Gilbert (2022) demonstrated how the cognitive offloading of setting external reminders may be driven by metamemory. They found that metamemory monitoring regions such as lateral PFC (lPFC), dorsal anterior cingulate cortex (dACC), and precuneus (McCurdy et al., 2013; Morales et al., 2018; Qiu et al., 2018; Ye et al., 2018) are involved in rating the confidence in remembering a delayed intention. These regions overlapped with the regions associated with generating a desire to use reminders (Boldt & Gilbert, 2022). In the present study, we built upon previous functional neuroimaging findings by utilizing diffusion tensor imaging (DTI), a white matter microstructural MRI technique, to explore whether the structural integrity of white matter tracts might be related to several features tested by metamemory driven cognitive offloading. We adapted a cognitive offloading task from Sachdeva and Gilbert (2020). In this task, participants perform a delayed intention task, either by being forced to use internal memory or external reminders, or freely choosing between the two strategies (Figure 1). Specifically, we focussed on the following three behavioral indices and their relationship with white matter tract structural integrity for this investigation:

**Fig. 1:**
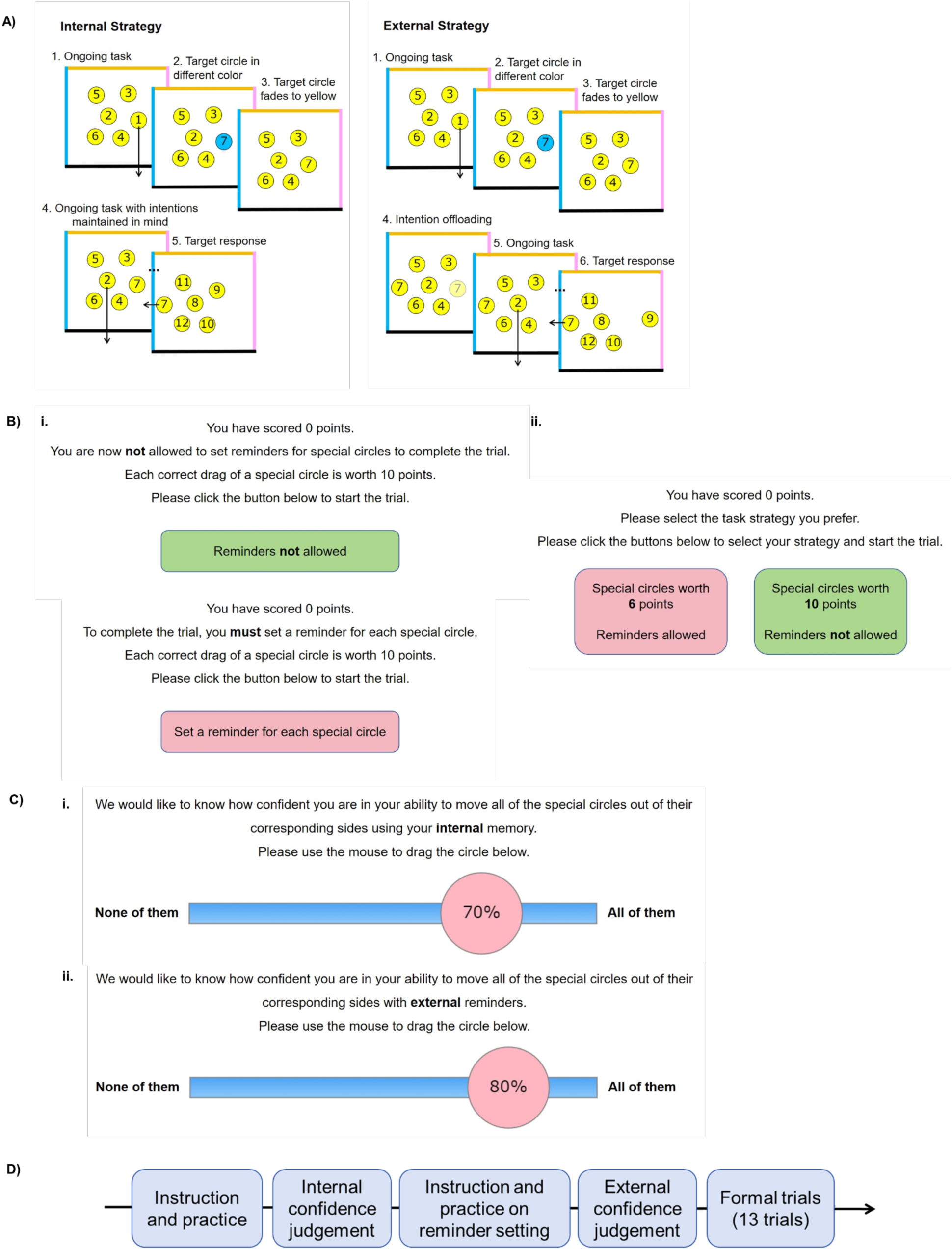
Task Procedure. **A)** In the delayed intention task, participants sequentially drag numbered circles (1–17) to the borders of a box. Most circles (10/17) are yellow and need placement at the bottom border. Occasionally, non-yellow circles (blue, orange, pink) appear, corresponding to the left, top, or right borders. These target circles must be dragged to their color-matched borders when their number in the sequence is reached. Non-yellow circles turn yellow after 2 seconds, requiring participants to form a delayed intention. Task points are accumulated for correct placements. In the internal strategy trials, participants use internal memory to complete the task. In the external strategy trials, participant can first drag the target circle near its corresponding border as a visual cue to remind them of the correct action. **B)** External reminder strategy selection: (i) In 3 out of 13 trials, participants were forced to rely on internal memory, and in another 3 trials, on external reminders. Correct actions earned 10 points regardless of strategy. (ii) In the remaining 7 trials, participants freely chose between strategies. External reminder actions earned 2–8 points, while internal memory actions always earned 10 points. **C)** Post-training confidence ratings: After practice trials, participants rated their confidence in their task performance using internal memory (i) or external reminders (ii). **D)** Task event sequence: An overview of the sequence of events in the task.

1) Confidence judgment: we focussed on participants’ own confidence rating (0–100%) regarding their prospective performance in trials where they had to rely solely on memory; 2) Metacognitive bias: the difference between one’s own confidence judgement and their actual performance (i.e., internal accuracy). Higher positive values reflect overconfidence, higher negative values reflect underconfidence, and values at zero indicate optimal calibration between confidence and performance; and 3) Reminder bias: the degree of preference for using external reminders over using memory in trials where participants could choose their strategy. Higher positive values indicate a strong bias toward external reminders, while higher negative values reflect a strong bias toward internal memory. In order to get a sense of how much the bias deviates from the optimal value irrespective of direction, we calculated the absolute values of metacognitive bias and reminder bias, denoted as |meta bias| and |reminder bias|, respectively.

Correspondingly, we selected a set of white matter fiber tracts, namely: Firstly, the superior longitudinal fasciculus (SLF). The SLF, which links the inferior parietal lobe to the lPFC, is known as a critical region for metacognitive monitoring and control (Qiu et al., 2018; Xue et al., 2023a; Zheng et al., 2021). We predicted that the higher structural integrity of the SLF to be associated with better calibration between confidence and accuracy, as well as a more optimal use of reminders. Secondly, the cingulum bundle (CB). Since the CB is anatomically connecting the precuneus, a well established metamemory related region (Ye et al., 2018) to dorsal anterior cingulate cortex (dACC), a cognitive control related region (Heilbronner & Haber, 2014; Shenhav et al., 2013), we hypothesized that the structural integrity of the CB would also be linked to better calibration between confidence and accuracy, and an optimal use of reminders. Thirdly, the fornix. Since the fornix is the afferent and efferent connecting fibre of the hippocampal formation to various subcortical structures that support episodic memory processes (Aggleton 2012; Kwok & Buckley, 2006, 2010; Kwok et al., 2015; Morand et al., 2021; Senova et al., 2020) and that higher-level metacognition must be built on the fundamental cognitive processes such as memory (Fleming & Daw, 2017; Nelson & Narens, 1990; Zheng et al., 2023), we hypothesized the structural integrity of the fornix to be related to the task performance when participants are forced to rely on internal memory, and also to the confidence prediction on their own task performance in this internal memory condition (Pan et al., 2024). Fourthly, the uncinate fasciculus (UF). Since the UF connects the temporal lobe including hippocampus to the rPFC and is known to support error monitoring in memory retrieval (Metzler-Baddeley et al., 2011; Petrides & Pandya, 2007), we hypothesized that higher structural integrity of the UF might lead to higher sensitivity in one own’s error and subsequently a more conservative confidence judgment.

## Results

### Behavioral results

We first investigated if there was a significant difference in accuracy between the forced internal memory and the forced external reminder conditions. We ran a 2×2 ANOVA and found that participants’ actual accuracy and confidence judgments were both higher when participants were forced to use external reminders than when they relied solely on internal memory (F(1, 128) = 41.21, p < .001; Figure 2A). Post-hoc tests confirmed that external reminders indeed increased the delayed intention task performance (*M* = 95.6583, *SD* = 8.4020) compared to the forced internal memory condition (*M* = 60.3642, *SD* = 15.54; *t*(50.775) = 11.649, *p* <.001). There was no main effect of measures (actual accuracy / self-judged accuracy, F(1, 128) = 0.059, p = .808) or interaction effect between types of conditions and measures (F(1, 128) = 0.61, p = .436), suggesting that participants’ confidence prediction matched their actual performance in both conditions.

**Figure 2.**
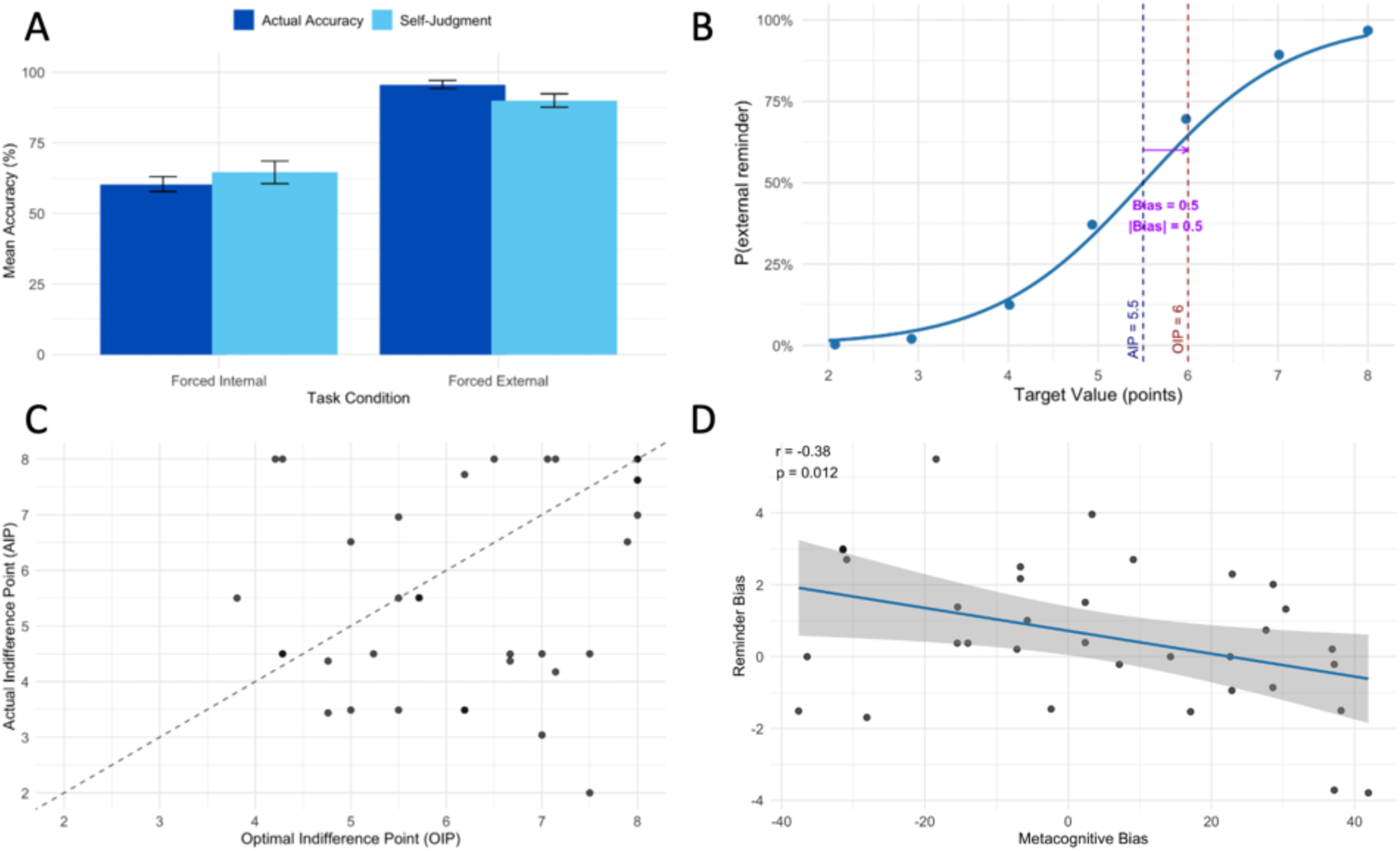
Metacognitive judgments of accuracy, computation of reminder bias using optimal indifference point (OIP) and actual indifference point (AIP), and relationship between metacognitive bias and reminder bias across participants. (A) Mean confidence judgments (self-reported accuracy estimates) and actual accuracy under forced-internal memory versus forced-external reminder conditions. (B) Illustration of reminder bias computation: the probability of choosing reminders across target values (2–8) is plotted with the optimal indifference point (OIP) and the actual indifference point (AIP). Reminder bias (shorten as |bias| in the plot) is quantified as the difference between OIP and AIP. (C) Scatterplot of all participants’ AIP versus OIP. The diagonal line denotes perfect calibration (AIP = OIP); points below it indicate over-reliance on reminders (AIP < OIP), points above indicate under-use of reminders (AIP > OIP). The group-level deviation from this line reveals a significant reminder bias. (D) Correlation plot of the relationship between reminder bias and metacognitive bias (confidence – actual accuracy), showing that participants who underestimate their memory accuracy (negative values on metacognitive bias) tend to rely more on reminders (positive values on reminder bias).

To quantify participants’ strategic preferences, we compared the *optimal indifference point* (OIP) and the *actual indifference point* (AIP). The OIP reflects the value at which an unbiased, reward-maximizing agent should be indifferent between using internal memory and external reminders. In contrast, the AIP denotes the actual value at which participants behaviorally exhibited indifference between the two strategies. With a one-sided paired t-test, we observed that the discrepancy between OIP and AIP (i.e., reminder bias, see illustration in Figure 2B) was significant, that is, the degree of reminder bias is significantly greater than zero (*M* = .5860, *SD* = 2.0357; *t*(33) = 1.6785, *p* = .0257). This statistical effect is reflected by the more dense individual dots below the diagonal line in Figure 2C. This pattern shows that the participants demonstrated a bias towards setting external reminders to assist their task performance, consistent with previous findings (e.g., Gilbert et al., 2020; Sachdeva & Gilbert, 2020).

Finally, a Pearson correlation test reveals a significantly negative correlation between reminder bias and the metacognitive bias for internal memory (*r*(32) = -.384, *p* = .0125; Figure 2D). This shows that participants who were under-confident in their own ability using internal memory were more likely to set up external reminders, confirming that the decision to use external reminders was a metacognition-guided strategy. All in all, at the behavioral level, our findings showed that the bias in setting up reminders is predicted by an estimate of one’s confidence towards subsequent performance in executing delayed intentions, consistent with previous reports (e.g., Gilbert et al., 2020).

### DTI Tractography and relationship with behavioral metrics

We then moved on to examine the relationship between these behavioral metrics and the participants’ white matter tract properties. Because reminder bias was driven by the discrepancy between self-estimated and actual performance under forced-internal memory, here we focused our analyses on metacognitive measures from the forced-internal memory condition. Given that the cognitive processes are often modulated by other factors, in all the following regression models, we input participants’ age, gender, and total intracranial volume as regressors of no interest to control for any potential influence on the regression results (Eikenes et al., 2023). Within each regression analysis, the outcome and all continuous predictors (i.e., except gender) were scaled first in order to better compare the coefficients across models.

We examined the relationship between the structure integrity of white matter (WM) tracts (indexed by fractional anisotropy; FA) and participants’ behavioral indices relevant to the metacognition driven cognitive offloading processes. FA reflects the microstructural integrity of white matter. High FA values suggest organized, healthy fiber tracts. The microstructural integrity within the superior longitudinal fasciculus (SLF), cingulum bundle (CB), and the fornix are implicated in relation to several aspects of participants’ performance in the cognitive offloading task. We did not obtain any significant results regarding the uncinate fasciculus (UF).

Firstly, the FA in the left and right SLF negatively predicted the absolute degree of bias (or deviation) in utilizing external reminders over internal memory (SLF L: ß = -0.437, *SE* = 0.159, t(29) = -2.753, *p* = 0.005, BF_10_ = 7.297; SLF R: ß = -0.351, *SE* = 0.178, t(29) = -1.972, *p* = 0.029, BF_10_ = 1.571, Table 1 & Figure 3A). These findings indicate that greater structural organization in the bilateral SLF was associated with a more optimal selection between external reminders and internal memory strategies.

**Table 1.**
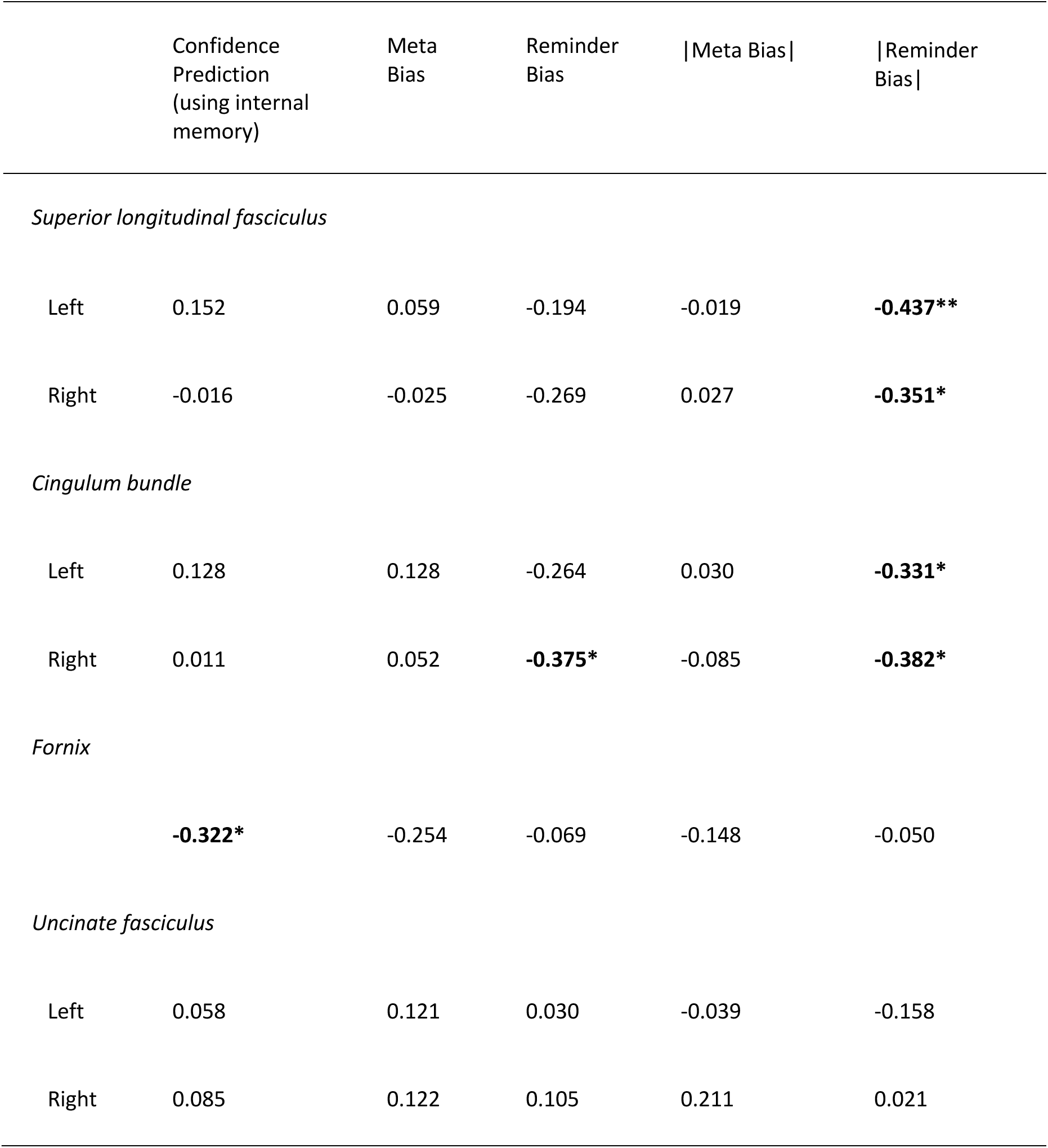
Regression coefficients between behavioral metrics and white matter tract FA index. Statistical significant coefficients are in bold, with ** indicating p-value < .01, * indicating p-value < .05, without family-wise error correction. Linear regression analyses were conducted where age, gender, and total intracranial volume on DTI metrics were controlled. Meta Bias = metacognitive bias. |Meta Bias| = absolute value of metacognitive bias; |Reminder Bias| = absolute value of reminder bias.

**Figure 3.**
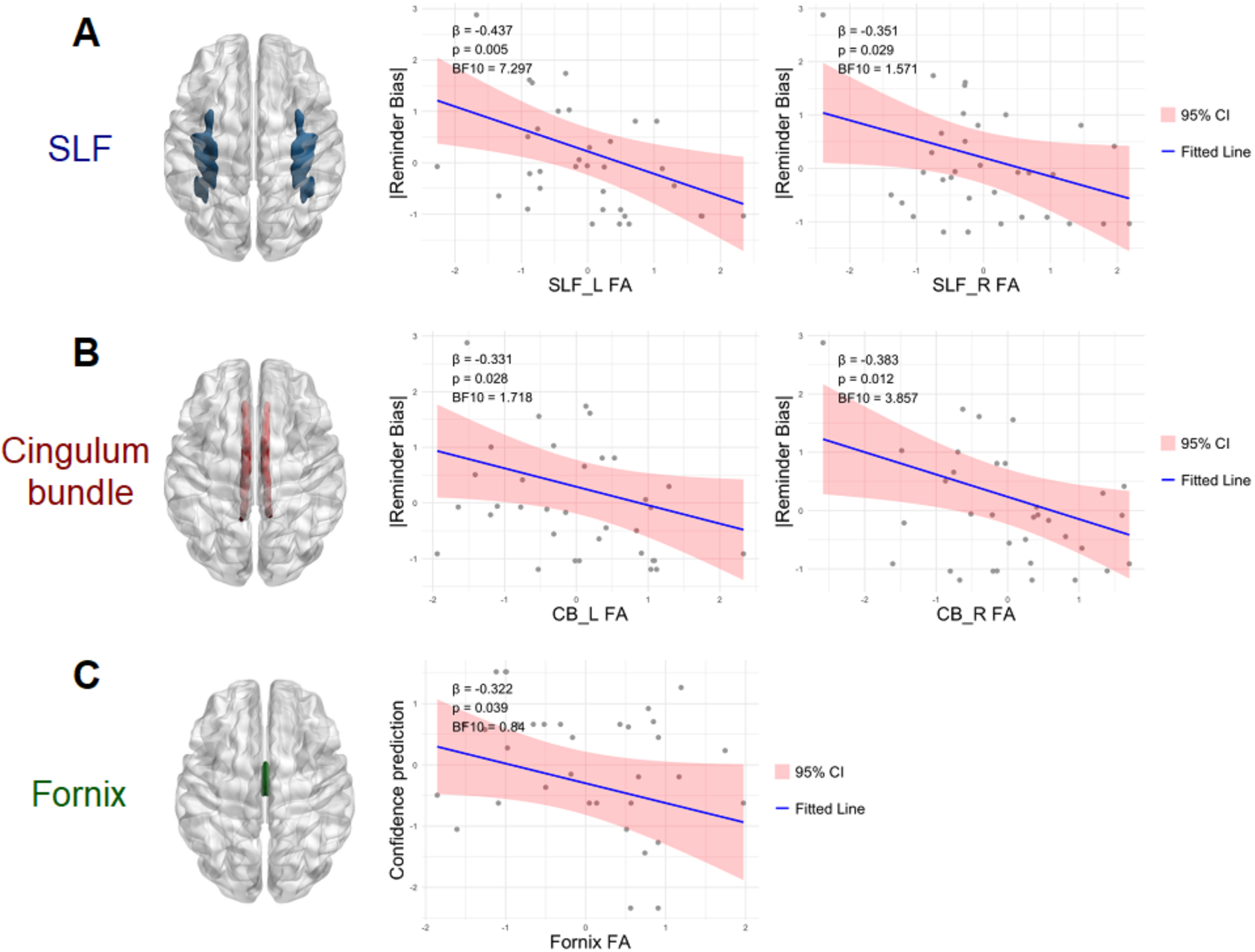
Correlations between behavior and DTI indices. A) Fractional anisotropy (FA) of the bilateral superior longitudinal fasciculus (SLF) was negatively associated with the absolute degree of deviation from the optimal strategy choice (|reminder bias|). B) Similarly, FA of the bilateral cingulum bundle (CB) negatively predicted the absolute bias in strategy selection (|reminder bias|). C) FA of the fornix was negatively associated with confidence levels when participants were required to rely solely on internal memory to perform delayed intention tasks. Regression coefficients (ß), P-values, and Bayes Factors are reported in top left within each regression plot. CI = 95% confidence interval. L = left hemisphere, R = right hemisphere.

Regarding the cingulum bundle, the microstructural integrity of the CB negatively also predicted the absolute degree of bias towards using external reminders over internal memory (CB L: ß = - 0.331, *SE* = 0.166, t(29) = -1.999, *p* = 0.028, BF_10_ = 1.718; CB R: ß = -0.383, *SE* = 0.160, t(29) = - 2.391, *p* = 0.012, BF_10_ = 3.857, Table 1 & Figure 3B). In other words, higher structural organization in the right CB corresponded to a more optimal strategy choice. Incidentally, the FA in the right CB also negatively predicted the degree of bias in setting up reminders ( ß = - 0.375, *SE* = 0.171, t(29) = -2.198, *p* = 0.018, BF_10_ = 1.906), which implies a reduced reliance on external reminders for those participants with more intact CB organization.

Thirdly, the microstructural integrity of the fornix negatively predicted the level of confidence in performing the offloading task with internal memory (ß = -0.322, *SE* = 0.176, t(29) = -1.827, *p* = 0.039, BF_10_ = 0.840; Table 1 & Figure 3C). This negative relationship implies, following the practice trials, participants with greater fornix microstructural organization adopt more conservative confidence predictions of their performance with respect to their internal memory.

We did not find any of the tracts to be associated with either metacognitive bias or the absolute magnitude of metacognitive bias. The regression results are summarized in Table 1.

## Discussion

In everyday life, people often use external reminders to offload memory loads and assist themselves in remembering delayed intentions. This cognitive offloading strategy is driven by a metacognitive sense of confidence (Dunn & Risko, 2016; Gilbert, 2023). When individuals are under-confident about their ability to execute delayed intentions, they are more likely to set up external reminders. Building on previous functional neuroimaging work (Boldt & Gilbert, 2022; Qiu et al., 2018), we utilized an anatomical neuroimaging technique, diffusion tensor imaging (DTI), to examine the relationship between white matter tract microstructural integrity and metrics embedded in metacognition-driven cognitive offloading. DTI maps the direction and magnitude of water diffusion in brain tissue, revealing the orientation and integrity of white matter tracts. Accordingly, we are able to use the data to infer anatomical connectivity—that is, how different brain regions are physically linked via axonal bundles.

At the white matter microstructural level, we found evidence highlighting how white matter structural organization in the superior longitudinal fasciculus (SLF) and cingulum bundle (CB) could predict deviations from optimal use of reminders. Anatomically, the SLF connects the lateral prefrontal cortex (lPFC) and the inferior parietal lobe (IPL). Prior studies have indicated that enhanced integrity of certain portions of the SLF is linked to better metacognitive monitoring across perceptual and mnemonic domains (Zheng et al., 2021). Similarly, stronger CB integrity negatively predicted the magnitude of bias toward reliance on reminders. People with weaker CB organizations tend to rely more excessively on external reminders. The CB links the dorsal anterior cingulate cortex (dACC) with parietal regions, including the precuneus, both of which are associated with metacognitive monitoring (Boldt & Gilbert, 2022; Morales et al., 2018; Rouault & Fleming, 2020; Ye et al., 2018). These findings imply that more efficient transmission of information due to the better microstructural organization through the SLF and CB seem to facilitate the use of metacognitive signals to guide offloading behavior. This aspect of cognitive offloading is considered as metacognitive control. Therefore, the current findings on these tracts, especially those on the SLF, extend on extant functional neuroimaging evidence that at the white matter microstructural level, monitoring and control processes might also share common anatomical and neural substrates (Boldt & Gilbert, 2022).

Compared to conventional metacognition research using trial-by-trial retrospective confidence ratings, here we asked participants to indicate their confidence prospectively on their task performance after the practice and training stages. This prospective confidence judgment can be interpreted as a global confidence judgment, which reflects broader, trait-like assessments of cognitive ability. Some studies suggest that global confidence judgments are derived from and constantly updated by trial-by-trial retrospective decision confidence (Lee et al., 2021; Mei et al., 2020; Rouault et al., 2019), which correspond to our practice and training stages. We found that structural organization of the fornix negatively predicted confidence predictions about one’s own subsequent memory task performance. Importantly, this relationship was not observed for actual memory performance or for the discrepancy between predicted and actual performance. A possible account is that participants with greater fornix integrity possess heightened sensitivity to errors committed during practice trials, via better hippocampal outputs to the ventral striatum (e.g., Aggleton, 2012). With richer error related information in the ventral striatum, a region known to encode prediction-error signals and to track global confidence on task performance (Rouault and Fleming, 2020), these participants may question their forthcoming task performance and thereby lower their confidence prediction. This interpretation is speculative, but our results may provide insights into the white matter microstructure underlying the global confidence judgments. Given that the fornix is an afferent and efferent of the hippocampal formation, the finding agrees with other recent work reporting that the hippocampus is implicated in mnemonic metacognition (Pan et al., 2024). In contrast with the fornix, SLF and CB primarily support transmission of the metacognitive signal and is related to dynamic, moment-to-moment metacognitive monitoring. This could explain why SLF and CB structural organization did not predict the level of confidence ratings nor metacognitive bias in our study.

Although the Bayesian power analysis indicates that a sample size of 34 participants provides sufficient power to detect medium effect sizes in a single multiple linear regression, it does not address the issue of multiple comparisons in the study. We acknowledge that the statistical power becomes more limited when conducting several regression analyses. The p-values we reported here should be interpreted with caution given the increased potential for false positives in the context of multiple comparisons. To draw more definitive conclusions, a large-scale replication will be necessary in the future. With a larger sample size, future research could investigate whether and how individual differences—such as age, gender, and psychiatric symptoms—influence metacognition driven cognitive offloading strategies through differences in brain microstructural properties (Boldt et al., 2025; Kirk et al., 2020; Scarampi & Gilbert, 2021; Tsai et al., 2022; Xue et al., 2023b). Incidentally, recent studies have shown that individuals possess a meta-metacognitive ability, namely, the capacity to evaluate the quality of their own confidence judgments (Recht et al., 2022; Zheng et al., 2023). Future work could therefore examine which white matter tracts also support meta-metacognition and how this capacity influences the optimal use of external reminders in prospective memory tasks.

In sum, our findings identify fronto-temporal-parietal white-matter tracts related to several metrics measured by metacognition driven cognitive offloading in a prospective memory task. These cognition-microstructural findings provide a link reflecting the relationship between cognitive offloading and the organization of white matter microstructures.

## Method and Materials

### Participants

38 adult participants were recruited through local advertisement. All participants had normal or corrected-to-normal vision, reported no history of no color blindness, psychiatric and neurological diseases, and no other contraindications for MRI. All participants provided written informed consent. The study was ethically approved by the Shanghai Jiao Tong University Institutional Review Board. All procedures were performed in accordance with the institutional guidelines.

Three participants were removed because they did not attend the MRI scanning. In line with the exclusion criteria used by Gilbert et al. (2020), we further excluded one participant who had a negative correlation between target value and likelihood of choosing to use reminders, implying a random or counter-rational strategy choice behavior. Therefore, 34 participants (19 females, mean age ± SD = 28.47 ± 8.82) were taken into analysis. Our Bayesian power analysis indicated that a sample size of 34 subjects provided high power to detect a medium effect size (regression coefficient of 0.5) in each of our planned subject-level multiple regression analyses.

### Task Procedure

In the following, we described three key stages of the task: Delayed intention, external reminder strategy selection, and pre-task confidence ratings.

#### Delayed intention

Participants engaged in a cognitive offloading task adapted from Gilbert et al. (2020). In this task, circles numbered from 1 to 17 would be presented within a box with colored borders. The primary objective for participants was to sequentially drag each circle to the bottom border in ascending numerical order (Fig. 1A). At the beginning of each trial, six yellow circles with numbers 1 to 6 were randomly positioned inside the box. Participants were instructed to drag these yellow circles to the bottom of the box. When a circle reached the box bottom, a new circle would replace it in its original location. For instance, dragging the circle numbered 1 to the bottom would be followed by the appearance of a circle numbered 7, taking its place.

Among the 17 circles per trial, most (10 out of 17) appeared in yellow and were to be dragged to the bottom edge as usual. However, on each trial, 7 of these circles initially appeared in a non-yellow color—blue, orange, or pink—corresponding to the left, top, or right edge of the box. These were designated as target circles. After 2 seconds, each colored target circle faded to yellow to match the others. Participants were instructed to remember both the digit and the original color of these target circles and to drag them to their color-matched edge when their digit was reached in the ongoing sequence. For example, after dragging Circle 1 to the bottom, an orange Circle 7 might appear and fade to yellow after 2 seconds; the participant must then continue dragging Circles 2–6 in order, while remembering to drag Circle 7 to the orange edge once it is reached.

This design required participants to maintain multiple delayed intentions — actually up to 6 at a time — throughout a sequence. This demand should be supraspan and exceed an individual’s typical short-term memory capacity. Moreover, we also randomized both the digits involved and target directions across trials, precluding fixed verbal encoding strategies (e.g., rote rehearsal like “digit seven-top, digit nine-left”). For illustration, we set up a link for a demo version of the task to help readers to try the task out for themselves: https://cognitiveoffloading.net/ZhengDemo. This is exactly the task from the real experiment we ran, except that it does not store any data and we have replaced all the Chinese instructions with the original English version from Sam Gilbert’s laboratory. For the overall structure of the paradigm, see Fig. 1D.

#### External reminder strategy selection

In addition to maintaining the delayed intention, participants were instructed how to set an external reminder for task performance. In such instances, participants needed to promptly drag a target circle near its corresponding box border upon appearance. For example, if a blue circle appeared, participants should immediately drag it close to the left border. This ensured that when the circle number was reached in the sequence, its location served as a reminder for participants to fulfill the intended action.

Each participant underwent 13 trials. In three trials, participants were explicitly instructed to rely solely on their internal representation of the intention, while in another set of three trials, participants were informed to rely exclusively on external reminders. In these forced-choice trials, every correct drag of a target circle resulted in a 10-point increase, regardless of the chosen strategy (Fig. 1B i). However, in the remaining seven trials, participants were given the freedom to choose their strategy (internal memory or external reminders) (Fig. 1B ii). Importantly, the points awarded for each correct drag varied based on the chosen strategy. Opting for internal memory yielded 10 points per accurate drag, while choosing external reminders resulted in a variable point range (2–8, randomly selected for each trial) for each correct drag, which was lower than the internal memory strategy. Therefore, simply always choosing to use reminders would not be an optimal strategy for task performance. Instead, participants have to balance the higher number of points when remembering with internal memory against the greater chance of success when using external reminders.

#### Post-training confidence ratings

Before the proper trials, participants underwent practice trials to familiarize themselves with the task and the two strategies. Following these practice trials, participants provided two separate confidence ratings (i.e., the prospective confidence) in task accuracy using internal memory (Fig. 1C i) or external reminders (Fig. 1C ii) respectively.

### Behavioral Indices

We used the following behavioral indices to quantify several aspects of each participant’s task performance. The first three are of particular interest in our present investigation in relation to the microstructural integrity of white matter tracts, whereas the next four were for replication of previous behavioral findings (see Boldt et al., 2024; Gilbert et al., 2020; Sachdeva & Gilbert, 2020) and to derive some of our key behavioral metrics.

> ***Reminder bias*** reflects one’s deviation from optimal reminder use. A value at zero indicates that a participant is optimal in selecting between internal memory and external reminder strategies. We also computed ***the absolute value of reminder bias*** (|Reminder Bias|) to capture the absolute magnitude of deviation from optimal reminder use. Mathematically, it is computed as the difference between Optimal indifference point (OIP) and Actual indifference point (AIP). An unbiased participant’s AIP should align with the OIP. Otherwise, a higher OIP than AIP indicates a bias toward external reminders because the participant would set up reminders even when the reward associated with the target is less than optimal. Conversely, a higher AIP than OPI would indicate a bias toward using internal memory (see below and FIgure 2B-C).

> ***Confidence judgment,*** participants’ own estimation on their likelihood that they can successfully move all the target circles by using internal memory only.

> ***Metacognitive bias***, the subtraction between confidence judgment and the objective percentage of targets remembered when using internal memory (i.e., the internal accuracy). The degree of metacognitive bias reflects how much a participant was over- or under-confident about their ability to remember targets with their internal memory. Negative values indicate under-confidence and vice versa, and a value at zero indicates that the participant is perfectly calibrated between their confidence prediction and actual task performance. We computed the ***absolute value of metacognitive bias*** (|Meta Bias|) to reflect the absolute magnitude of miscalibration in confidence judgments.

The following four were not of our main focus but were used to derive the three aforementioned metrics and for replication of previous behavioral findings.

> ***Internal accuracy*** (ACC_FI_), the average percentage of target circles correctly moved to designated locations in trials where participants were required to rely solely on internal memory (i.e., forced internal trials).

> ***External accuracy*** (ACC_FE_), the average percentage of target circles correctly moved to designated locations in trials where participants were required to rely solely on external reminders (i.e., forced external trials).

> ***Optimal indifference point*** (OIP), the value of target circles at which an unbiased, reward-maximizing participant should be indifferent towards either the internal strategy or external strategy. OIP is calculated as 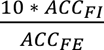. Suppose a participant can accurately remember 60% of targets with internal memory (ACC_FI_ = 0.6) and all targets with external reminders (ACC_FI_ = 1), then in this case the OIP value should be 6. That is, obtaining 6 points per target with external reminders would lead to the same amount of points as obtaining 10 points per target with internal memory. When the value of a target circle is above 6, it would be optimal to choose external reminders, as it maximizes the rewards. Similarly, the value of a target circle is below 6, it would be optimal to choose internal memory.

> ***Actual indifference point*** (AIP), the actual value of target circles at which participants showed indifference. The AIP is determined by assessing the probability of choosing an external strategy over an internal one across all external target values, and was obtained by fitting a psychometric function using the R package “*quickspy*.”

### DTI image acquisition

All participants underwent magnetic resonance imaging (MRI) on a 3-tesla system equipped with a standard 32-channel head and neck coil (uMR790, United Imaging, Shanghai, China). The MRI protocol included the following sequences: (1) 3D T_1_-weighted gradient-recalled echo sequence with repetition time (TR) = 8.1 ms, echo time (TE) = 3.4 ms, inversion time (TI) = 1060 ms, matrix size = 320 × 300 × 208, voxel size = 0.8 mm isotropic, flip angle = 8°, bandwidth = 260 Hz/pixel, acceleration factor = 2, and field of view (FOV) = 256 × 240 mm². (2) Multi-shell diffusion-weighted imaging (DWI) using multi-band accelerated echo planar imaging (EPI): 96 diffusion-weighted directions, including 32 directions at b = 1000 s/mm² and 64 directions at b = 2000 s/mm², along with 4 b = 0 images. The acquisition parameters for the DWI included: voxel size = 1.5 mm, TR = 5150 ms, TE = 77 ms, FOV = 210 × 210 mm², and flip angle = 90°.

### DTI data processing

The multi-shell diffusion magnetic resonance imaging (dMRI) data underwent preprocessing for denoising and removal of Gibbs artifacts utilizing tools from MRtrix 3.0 (http://www.mrtrix.org). Following this, head-motion correction and eddy current correction were carried out using the Functional Magnetic Resonance Imaging of the Brain Software Library (FSL; v6.0; https://fsl.fmrib.ox.ac.uk/fsl/fslwiki). Subsequently, the diffusion tensor fitting for all diffusion shells with b-values up to 1000 s/mm² was performed using the *dtifit* function within FSL. The fractional anisotropy (FA) images were obtained for each participant.

White matter tracts of interest—including the column and body of the fornix, bilateral uncinate fasciculus (UF), cingulum bundle, and superior longitudinal fasciculus (SLF)—were delineated based on the ICBM-DTI-81 white matter labels atlas. To accurately register these masks onto the FA maps, diffusion-weighted images were first co-registered to T1-weighted structural images and subsequently normalized to Montreal Neurological Institute (MNI) space using the transformation matrix obtained from structural image normalization. The inverse of these transformations was then applied to the masks to align them within the native diffusion-weighted image space. All registration procedures were executed using Advanced Normalization Tools (ANTs, http://stnva.github.io/ANTs/). To mitigate potential registration inaccuracies and minimize partial volume effects with cerebrospinal fluid (CSF), the registered masks were further refined by intersecting them with each subject’s white matter (WM) mask. The mean values of FA were then computed for each subject’s fornix, UF, cingulum bundle, and SLF, incorporating all voxels within the respective tracts. The FA is interpreted as a quantitative biomarker of white matter ‘integrity’ (for reviews see Assaf and Pasternak, 2008; Beaulieu, 2009). We will use this biomarker for our investigation of cognitive processes.

#### Statistical analysis and code availability

All the pairwise t-tests and linear regression analyses in this study were conducted using the R “stats” package. In the regression analyses, all the continuous outcome and predictors were firstly scaled, in order to better compare the coefficients across models. We used the R “BayesFactor” package for the Bayes factor computation for each statistical analysis, as well as for the Bayesian power analysis to determine whether the present study has sufficient power to detect small to medium effect sizes. The preprocessed behavioral and DTI data, along with all relevant analysis code, are available at https://osf.io/jdnk4/.

## Author Contributions

Conceptualization, YZ, SG, SCK, YL ; methodology, YZ, SG, SCK, YL; investigation, YZ, BB, DW, SCK, YL ; formal analysis, YZ, BB, DW, YyL, SG, SCK, YL; visualization, YZ, BB, YyL, DW, YL.; writing – original draft, YZ, BB, YyL.; writing – review & editing, YZ, SG, SCK, YL; supervision, SCK, YL ; funding acquisition, SCK, YL.

## Data availability

Data and materials are available at https://osf.io/jdnk4/.

## Acknowledgements

This work received support from Jiangsu Provincial Department of Science and Technology (BK20221267) and an internal funding from School of Psychology and Cognitive Science (East China Normal University) to SCK; Shanghai Pilot Program for Basic Research-Shanghai Jiao Tong University (No. 21TQ1400203), Program for Professor of Special Appointment (Eastern Scholar) at Shanghai Institutions of Higher Learning to YL.

## References

Assaf, Y., & Pasternak, O. (2008). Diffusion tensor imaging (DTI)-based white matter mapping in brain research: a review. Journal of Molecular Neuroscience, 34, 51–61.

Aggleton, J. P. (2012). Multiple anatomical systems embedded within the primate medial temporal lobe: implications for hippocampal function. Neuroscience & Biobehavioral Reviews, 36(7), 1579–1596.

Beaulieu, C., 2009. The biological basis of diffusion anisotropy. In: Johansen-Berg, H., Behrens, T.E.J. (Eds.), Diffusion MRI: From Quantitative Measurement to In-vivo Neuroanatomy. Academic Press, pp. 105–126.

Boldt, A., Fox, C. A., Gillan, C. M., & Gilbert, S. (2025). Transdiagnostic compulsivity is associated with reduced reminder setting, only partially attributable to overconfidence. eLife, 13.

Boldt, A., & Gilbert, S. J. (2022). Partially Overlapping Neural Correlates of Metacognitive Monitoring and Metacognitive Control. The Journal of Neuroscience, 42(17), 3622–3635. 10.1523/JNEUROSCI.1326-21.2022

Boldt, A., Schiffer, A.-M., Waszak, F., & Yeung, N. (2019). Confidence Predictions Affect Performance Confidence and Neural Preparation in Perceptual Decision Making. Scientific Reports, 9(1), Article 1. 10.1038/s41598-019-40681-9

Burgess, P. W., Quayle, A., & Frith, C. D. (2001). Brain regions involved in prospective memory as determined by positron emission tomography. Neuropsychologia, 39(6), 545–555. 10.1016/S0028-3932(00)00149-4

Burgess, P. W., Scott, S. K., & Frith, C. D. (2003). The role of the rostral frontal cortex (area 10) in prospective memory: A lateral versus medial dissociation. Neuropsychologia, 41(8), 906–918. 10.1016/S0028-3932(02)00327-5

Chiu, G., & Gilbert, S. J. (2024). Influence of the physical effort of reminder-setting on strategic offloading of delayed intentions. Quarterly Journal of Experimental Psychology, 77(6), 1295– 1311. 10.1177/17470218231199977

Crovitz, H. F., & Daniel, W. F. (1984). Measurements of everyday memory: Toward the prevention of forgetting. Bulletin of the Psychonomic Society, 22(5), 413–414. 10.3758/BF03333861

Dunn, T. L., & Risko, E. F. (2016). Toward a Metacognitive Account of Cognitive Offloading. Cognitive Science, 40(5), 1080–1127. 10.1111/cogs.12273

Dupont, D., Zhu, Q., & Gilbert, S. J. (2023). Value-based routing of delayed intentions into brain-based versus external memory stores. Journal of Experimental Psychology: General, 152, 175–187. 10.1037/xge0001261

Eichenbaum, H. (2017). Prefrontal–hippocampal interactions in episodic memory. Nature Reviews Neuroscience, 18(9), 547–558. 10.1038/nrn.2017.74

Eikenes, L., Visser, E., Vangberg, T., & Håberg, A. K. (2023). Both brain size and biological sex contribute to variation in white matter microstructure in middle-aged healthy adults. Human Brain Mapping, 44(2), 691–709.

Eskritt, M., & Ma, S. (2014). Intentional forgetting: Note-taking as a naturalistic example. Memory & Cognition, 42(2), 237–246. 10.3758/s13421-013-0362-1

Frömer, R., Lin, H., Dean Wolf, C. K., Inzlicht, M., & Shenhav, A. (2021). Expectations of reward and efficacy guide cognitive control allocation. Nature Communications, 12(1), Article 1. 10.1038/s41467-021-21315-z

Gilbert, S. J. (2011). Decoding the Content of Delayed Intentions. The Journal of Neuroscience, 31(8), 2888–2894. 10.1523/JNEUROSCI.5336-10.2011

Gilbert, S. J. (2015). Strategic use of reminders: Influence of both domain-general and task-specific metacognitive confidence, independent of objective memory ability. Consciousness and Cognition, 33, 245–260. 10.1016/j.concog.2015.01.006

Gilbert, S. J. (2023). Cognitive offloading is value-based decision making: Modelling cognitive effort and the expected value of memory. PsyArXiv. 10.31234/osf.io/5e8mg

Gilbert, S. J. (2024). Cognitive offloading is value-based decision making: Modelling cognitive effort and the expected value of memory. Cognition, 247, 105783. 10.1016/j.cognition.2024.105783

Gilbert, S. J., Bird, A., Carpenter, J. M., Fleming, S. M., Sachdeva, C., & Tsai, P.-C. (2020). Optimal use of reminders: Metacognition, effort, and cognitive offloading. Journal of Experimental Psychology: General, 149, 501–517. 10.1037/xge0000652

Gilbert, S. J., Boldt, A., Sachdeva, C., Scarampi, C., & Tsai, P.-C. (2023). Outsourcing Memory to External Tools: A Review of ‘Intention Offloading.’ Psychonomic Bulletin & Review, 30(1), 60–76. 10.3758/s13423-022-02139-4

Heilbronner, S. R., & Haber, S. N. (2014). Frontal Cortical and Subcortical Projections Provide a Basis for Segmenting the Cingulum Bundle: Implications for Neuroimaging and Psychiatric Disorders. Journal of Neuroscience, 34(30), 10041–10054. 10.1523/JNEUROSCI.5459-13.2014

Hu, X., Luo, L., & Fleming, S. M. (2019). A role for metamemory in cognitive offloading. Cognition, 193, 104012. 10.1016/j.cognition.2019.104012

Kelly, M. O., & Risko, E. F. (2019). The isolation effect when offloading memory. Journal of Applied Research in Memory and Cognition, 8, 471–480. 10.1037/h0101842

Kelly, M. O., & Risko, E. F. (2022). Study effort and the memory cost of external store availability. Cognition, 228, 105228. 10.1016/j.cognition.2022.105228

Kwok, S. C., & Buckley, M. J. (2006). Fornix transection impairs exploration but not locomotion in ambulatory macaque monkeys. Hippocampus, 16(8), 655–663. 10.1002/hipo.20195

Kwok, S. C., & Buckley, M. J. (2010). Fornix transection selectively impairs fast learning of conditional visuospatial discriminations. Hippocampus, 20(3), 413–422. 10.1002/hipo.20643

Kwok, S. C., Mitchell, A. S., & Buckley, M. J. (2015). Adaptability to changes in temporal structure is fornix-dependent. Learning & Memory, 22(8), 354–359. 10.1101/lm.038851.115

Landsiedel, J., & Gilbert, S. J. (2015). Creating external reminders for delayed intentions: Dissociable influence on “task-positive” and “task-negative” brain networks. NeuroImage, 104, 231–240.10.1016/j.neuroimage.2014.10.021

Lee, A. L. F., de Gardelle, V., & Mamassian, P. (2021). Global visual confidence. Psychonomic Bulletin & Review, 28(4), 1233–1242. 10.3758/s13423-020-01869-7

McCurdy, L. Y., Maniscalco, B., Metcalfe, J., Liu, K. Y., Lange, F. P. de, & Lau, H. (2013). Anatomical Coupling between Distinct Metacognitive Systems for Memory and Visual Perception. Journal of Neuroscience, 33(5), 1897–1906. 10.1523/JNEUROSCI.1890-12.2013

Mei, N., Rankine, S., Olafsson, E., & Soto, D. (2020). Similar history biases for distinct prospective decisions of self-performance. Scientific Reports, 10(1), 5854. 10.1038/s41598-020-62719-z

Metzler-Baddeley, C., Jones, D. K., Belaroussi, B., Aggleton, J. P., & O’Sullivan, M. J. (2011). Frontotemporal Connections in Episodic Memory and Aging: A Diffusion MRI Tractography Study. The Journal of Neuroscience, 31(37), 13236–13245. 10.1523/JNEUROSCI.2317-11.2011

Momennejad, I., & Haynes, J.-D. (2013). Encoding of Prospective Tasks in the Human Prefrontal Cortex under Varying Task Loads. Journal of Neuroscience, 33(44), 17342–17349. 10.1523/JNEUROSCI.0492-13.2013

Morales, J., Lau, H., & Fleming, S. M. (2018). Domain-General and Domain-Specific Patterns of Activity Supporting Metacognition in Human Prefrontal Cortex. The Journal of Neuroscience: The Official Journal of the Society for Neuroscience, 38(14), 3534–3546. 10.1523/JNEUROSCI.2360-17.2018

Morand, A., Segobin, S., Lecouvey, G., Gonneaud, J., Eustache, F., Rauchs, G., & Desgranges, B. (2021). Brain Substrates of Time-Based Prospective Memory Decline in Aging: A Voxel-Based Morphometry and Diffusion Tensor Imaging Study. Cerebral Cortex, 31(1), 396–409. 10.1093/cercor/bhaa232

Pan, K., Guo, X., Zheng, Z., Zhu, Z., Wu, H., Zhu, J., … & Jiang, H. (2024). Components of Mnemonic Metacognition Constitutionally Supported by High Gamma Activity Between the Precuneus and Hippocampus. bioRxiv, 10.1101/2024.02.26.581526

Petrides, M., & Pandya, D. N. (2007). Efferent Association Pathways from the Rostral Prefrontal Cortex in the Macaque Monkey. The Journal of Neuroscience, 27(43), 11573–11586. 10.1523/JNEUROSCI.2419-07.2007

Qiu, L., Su, J., Ni, Y., Bai, Y., Zhang, X., Li, X., & Wan, X. (2018). The neural system of metacognition accompanying decision-making in the prefrontal cortex. PLOS Biology, 16(4), e2004037. 10.1371/journal.pbio.2004037

Recht, S., Jovanovic, L., Mamassian, P., & Balsdon, T. (2022). Confidence at the limits of human nested cognition. Neuroscience of Consciousness, 2022(1), niac014.

Risko, E. F., & Dunn, T. L. (2015). Storing information in-the-world: Metacognition and cognitive offloading in a short-term memory task. Consciousness and Cognition, 36, 61–74. 10.1016/j.concog.2015.05.014

Risko, E. F., & Gilbert, S. J. (2016). Cognitive Offloading. Trends in Cognitive Sciences, 20(9), 676–688. 10.1016/j.tics.2016.07.002

Rouault, M., Dayan, P., & Fleming, S. M. (2019). Forming global estimates of self-performance from local confidence. Nature Communications, 10, 1141. 10.1038/s41467-019-09075-3

Rouault, M., & Fleming, S. M. (2020). Formation of global self-beliefs in the human brain. Proceedings of the National Academy of Sciences, 117(44), 27268–27276. 10.1073/pnas.2003094117

Rummel, J., & Kvavilashvili, L. (2023). Current theories of prospective memory and new directions for theory development. Nature Reviews Psychology, 2(1), 40–54. 10.1038/s44159-022-00121-4

Sachdeva, C., & Gilbert, S. J. (2020). Excessive use of reminders: Metacognition and effort-minimisation in cognitive offloading. Consciousness and Cognition, 85, 103024. 10.1016/j.concog.2020.103024

Scarampi, C., & Gilbert, S. J. (2021). Age differences in strategic reminder setting and the compensatory role of metacognition. Psychology and Aging, 36(2), 172–185.

Senova, S., Fomenko, A., Gondard, E., & Lozano, A. M. (2020). Anatomy and function of the fornix in the context of its potential as a therapeutic target. *Journal of Neurology*, Neurosurgery & Psychiatry, 91(5), 547–559. 10.1136/jnnp-2019-322375

Shenhav, A., Botvinick, M. M., & Cohen, J. D. (2013). The Expected Value of Control: An Integrative Theory of Anterior Cingulate Cortex Function. Neuron, 79(2), 217–240. 10.1016/j.neuron.2013.07.007

Simons, J. S., Schölvinck, M. L., Gilbert, S. J., Frith, C. D., & Burgess, P. W. (2006). Differential components of prospective memory?: Evidence from fMRI. Neuropsychologia, 44(8), 1388– 1397. 10.1016/j.neuropsychologia.2006.01.005

Sparrow, B., Liu, J., & Wegner, D. M. (2011). Google Effects on Memory: Cognitive Consequences of Having Information at Our Fingertips. Science, 333(6043), 776–778. 10.1126/science.1207745

Tsai, P. C., Scarampi, C., Kliegel, M., & Gilbert, S. J. (2023). Optimal cognitive offloading: Increased reminder usage but reduced proreminder bias in older adults. Psychology and Aging, 38(7), 684–695.

Xue, K., Zheng, Y., Rafiei, F., & Rahnev, D. (2023a). The timing of confidence computations in human prefrontal cortex. Cortex, 168, 167–175. 10.1016/j.cortex.2023.08.009

Xue, K., Zheng, Y., Papalexandrou, C., Hoogervorst, K., Allen, M., & Rahnev, D. (2023b). *No Gender Difference in Confidence or Metacognitive Ability in Perceptual Decision Making* (SSRN Scholarly Paper 4820298). 10.2139/ssrn.4820298

Ye, Q., Zou, F., Lau, H., Hu, Y., & Kwok, S. C. (2018). Causal Evidence for Mnemonic Metacognition in Human Precuneus. The Journal of Neuroscience, 38(28), 6379–6387. 10.1523/JNEUROSCI.0660-18.2018

Zheng, Y., Recht, S., & Rahnev, D. (2023). Common computations for metacognition and meta-metacognition. Neuroscience of Consciousness, 2023(1), niad023. 10.1093/nc/niad023

Zheng, Y., Wang, D., Ye, Q., Zou, F., Li, Y., & Kwok, S. C. (2021). Diffusion property and functional connectivity of superior longitudinal fasciculus underpin human metacognition. Neuropsychologia, 156, 107847. 10.1016/j.neuropsychologia.2021.107847

